# Understanding policing as a mechanism of cheater control in cooperating bacteria

**DOI:** 10.1101/267799

**Authors:** Tobias Wechsler, Rolf Kümmerli, Akos Dobay

**Author notes:** **Email addresses** (Akos Dobay). **Corresponding author**: Akos Dobay, Zurich Institute of Forensic Medicine, University of Zurich, Winterthurerstrasse 190/52, CH-8057 Zurich, Switzerland.

## Abstract

Policing occurs in insect, animal and human societies, where it is used as a conditional strategy to prevent cheating and enforce cooperation. Recently, it has been suggested that policing might even be relevant in enforcing cooperation in much simpler organisms such as bacteria. Here, we used individual-based modelling to develop an evolutionary concept for policing in bacteria, and identify the conditions under which it can be adaptive. We modelled interactions between cooperators, producing a beneficial public good, cheaters exploiting the public good without contributing to it, and public good producing policers that secrete a toxin to selectively target cheaters. We found that toxin-mediated policing is favored when (i) toxins are potent and durable, (ii) cheap to produce, (iii) cell and public good diffusion is intermediate, and (iv) toxins diffuse farther than the public good. Overall, we show that toxin-mediated policing can enforce cooperation, but the parameter space where it is beneficial seems quite narrow. Moreover, we found that policing decays when the genetic linkage between public good and toxin production breaks. This is because policing is itself a public good, offering protection to toxin-resistant mutants that still produce public goods, yet no longer invest in toxins. Our work suggests that very specific environmental conditions are required for genetically fixed policing mechanisms to evolve in bacteria, and offers empirically testable predictions for their evolutionary stability.

## Introduction

Cooperation, a process where individuals act to increase the fitness of others at an immediate cost to themselves, is common across the tree of life (Sachs et al. 2004; West et al. 2007b). While cooperation is widespread in humans (Fehr and Fischbacher 2003) and higher organisms such as insects and vertebrates (Clutton-Brock 2002; Ratnieks et al. 2006), we have only recently recognized that microbes also evolved adaptive cooperative behavior (West et al. 2007a; Nadell et al. 2009; Ross-Gillespie and Kümmerli 2014; Cavaliere et al. 2017). Types of microbial cooperation include the formation of biofilms, where individuals secrete polymeric compounds to build a protective extracellular matrix (Nadell et al. 2009), the formation of fruiting bodies where some individuals sacrifice themselves to enable the dispersal of others (Velicer and Vos 2009); and the secretion of shareable metabolites, including proteases to digest nutrients (Diggle et al. 2007; Sandoz et al. 2007), siderophores to scavenge extra-cellular iron (Griffin et al. 2004; Kümmerli et al. 2010), and biosurfactants enabling group motility (de Vargas Roditi et al. 2013).

Although cooperation is thought to provide benefits for the collective as a whole, it is intrinsically vulnerable to exploitation by cheating mutants, which benefit from the cooperative acts performed by others, but refrain from contributing themselves to the welfare of the group (Ghoul et al. 2014). Thus, there has been great interest in identifying mechanisms that maintain cooperation and prevent the invasion of cheaters (West et al. 2007b; Asfahl and Schuster 2017; Özkaya et al. 2017). Work on microbes have proved particularly useful in this context because cheating mutants can easily be engineered and factors important for cooperation can be experimentally manipulated in laboratory settings. Studies following this approach revealed a plethora of mechanisms promoting cooperation, including: (i) limited dispersal ensuring that cooperators stay together (Kümmerli et al. 2009; van Gestel et al. 2014); (ii) molecular mechanisms allowing the recognition of other cooperators (Mehdiabadi et al. 2006; Smukalla et al. 2008; Rendueles et al. 2015); (iii) regulatory linkage between multiple traits imposing additional costs on cheaters (Jousset et al. 2009; Dandekar et al. 2012; Ross-Gillespie et al. 2015; Mitri and Foster 2016; Popat et al. 2017); and (iv) mechanisms reducing the cost of cooperation such as public good recycling (Kümmerli and Brown 2010) or the use of superfluous nutrients for public goods production (Xavier et al. 2011; Sexton and Schuster 2017).

In addition, several studies suggested that bacteria have evolved policing mechanisms, which enable cooperators to directly sanction cheaters and thereby enforce cooperation (Manhes and Velicer 2011; Inglis et al. 2014; Wang et al. 2015; Majerczyk et al. 2016; Evans et al. 2018). For example, Wang et al. (2015) showed that in the opportunistic pathogen *Pseudomonas aeruginosa*, the synthesis of a publicly shareable protease is regulatorily coupled to the synthesis of the toxin cyanide in such a way that cheaters, deficient for protease production, simultaneously lose immunity against cyanide. Thus, the costly synthesis of a harmful toxin, which selectively targets non-cooperative cheaters, can be understood as a policing mechanism. While it seems intriguing that organisms as simple as bacteria can perform policing behavior, multiple questions regarding the evolution of microbial policing have remained unaddressed. For one thing, we know little about the environmental conditions promoting policing via toxin production. In this context, one would expect that the spatial structure of the environment, which affects the diffusion of cells, public goods and toxins, should play a crucial role (Driscoll and Pepper 2010; Inglis et al. 2011; Dobay et al. 2014). Moreover, it is unknown how potent a toxin must be and how much it can cost in order to efficiently fight cheaters. Finally, it remains unclear whether policing is an evolutionary stable strategy or whether non-policing cooperators, which are immune against toxins can exploit policers as second order cheaters (Inglis et al. 2014), as it is the case in higher organisms (Boyd et al. 2003).

Here, we address these issues by using realistic individual-based models that simulate interactions between cooperating, policing and cheating bacteria on a toroidal two-dimensional surface (Dobay et al. 2014). In our *in-silico* approach, bacteria are modeled as discs and seeded in low numbers onto the surface of their habitat, where they can consume resources, grow, divide, disperse, and secrete compounds according to specified parameters. Important to note is that both bacteria and public good molecules are modelled as individual agents and are free to diffuse on a continuous landscape, closely mimicking biological conditions (Figs. 1 + S1). For our main analysis, we considered two bacterial strains that match those used in the empirical policing paper by Wang et al. (2015). Specifically, we implemented a policing strain *P* that produces a public good together with a tightly linked toxin and an immunity protein; and a cheating strain *C* that is deficient for public good production and, due to the regulatory linkage, is also unable to produce the toxin and its immunity protein. We further considered a cooperating control wildtype strain *W*, which produces the public good but features no policing mechanism. Finally, we also considered a second-order policing cheater *R*, which produces the public good and the immunity protein, but not the toxin. For this strain, we assume that the genetic linkage between the three traits can be broken, such that *R* directly evolves from *P*.

**Figure 1.**
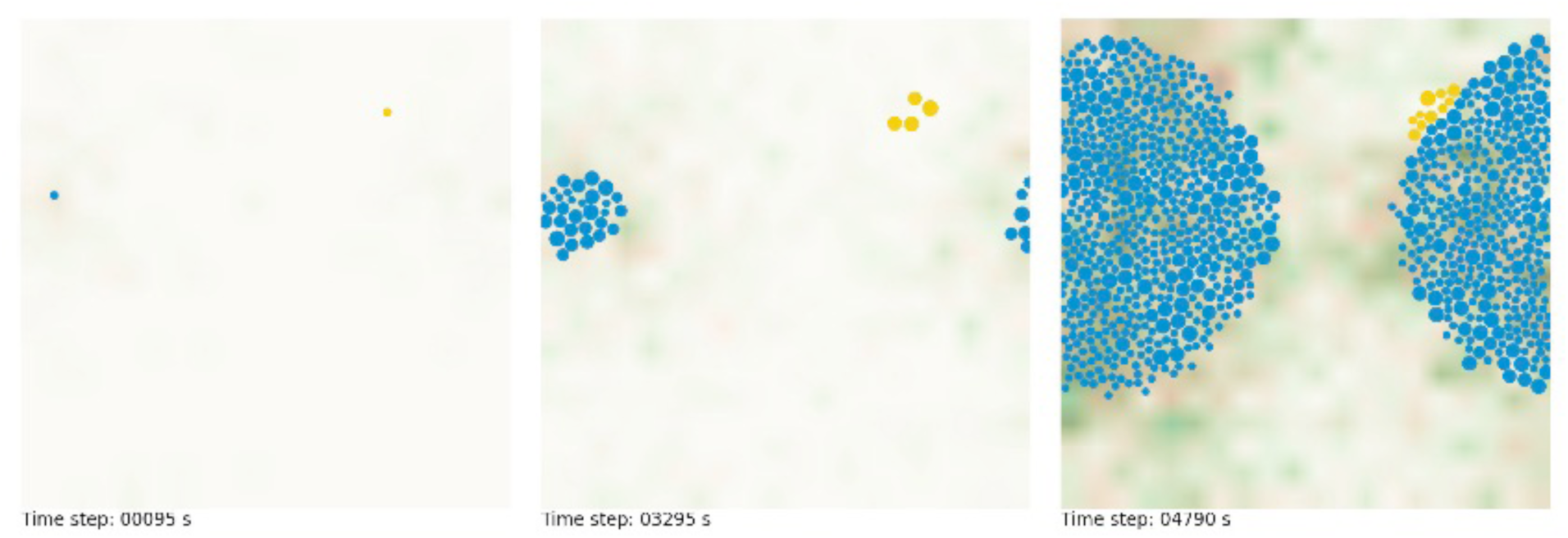
Point-in-time snapshots showing the spatial distribution of cooperating policers (in blue) and cheaters (in yellow) during competition. The point-in-time snapshots were taken during the early, intermediate and late simulation phase, depicting the growth of digital bacteria in their simulated environment. The density of public goods (in green) and toxins (in red) produced by the cooperating policers are indicated by the intensity level of their respective color. The specific parameters values for this example are: *D*_c_ = 0.0 µm^2^/s, *d*_pg_ = 1.0 µm^2^/s, *d*_tox_ = 2.0 µm^2^/s, *δ*_*tox*_ = 500 s, *θ*_*T*_= 1000 and κ = 3.5.

In a first set of simulations, we examined the performance of the cooperator, cheater and policing strains in monoculture to understand how the relative costs and benefits of public good and toxin production affect strain fitness. Next, we competed wildtype cooperators against cheaters across a range of bacterial and public good diffusion coefficients to determine the parameter space where cooperation is favored in the absence of policing (Dobay et al. 2014). Subsequently, we competed the cheater against the policing strain across the same parameter space, to test whether policing extends the range of conditions across which cooperation is favored, and to identify properties of the toxin system (diffusion, potency, durability) to guarantee efficient policing. Finally, we simulated the situation where toxin-resistant public good producers evolve from the policing strain, and ask whether policing via toxin production itself constitutes an exploitable public good.

## Methods

### *IN-SILICO* HABITAT AND BACTERIA

The *in-silico* habitat consists of a two-dimensional continuous toroidal surface, with no boundaries. The size of the surface is 60 x 60 µm^2^ = 3,600 µm^2^. Bacteria are modeled as discs with an initial radius *r* of 0.5 µm. Bacteria can consume resources, which are unlimited in our system, grow at a basic growth rate µ, and divide when reaching the threshold radius of 1.0 µm. Bacteria can disperse on the landscape according to a specific cell diffusion coefficient *D*_c_ (µm^2^/s), and are not bound to a grid, but can freely move on the continuous landscape (i.e. we used an off-lattice model with double-precision floating-point format). At the beginning of a simulation, we randomly seed one founder bacteria of each strain onto the surface, free to grow and divide according to its life cycle (see Figs. 1 + S1 for a visualization).

In addition to the basic growth rate µ, the growth of a bacterium is influenced by its social behavior and its interaction with other community members. Costs, reducing growth, incur to individuals producing public goods, toxins and immunity proteins. Additional costs incur to susceptible cells taking up toxins. The uptake of a public good, meanwhile, has a positive effect on growth for the beneficiary. Accordingly, the growth of our four strains *W* (public good producing wildtype strain), *P* (policing strain), *C* (toxin-sensitive cheating strain), and *R* (second-order toxin-resistant cheater strain) are defined by the following set of functions:

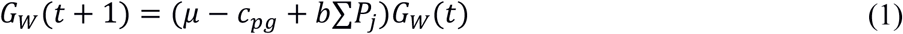

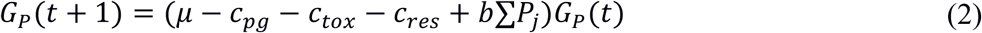

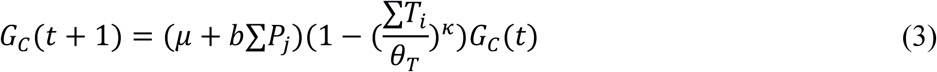

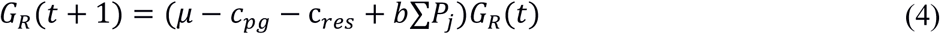

where *c*_*pg*_, *c*_*tox*_, *c*_*res*_ are the production costs per public good, toxin and the immunity protein, respectively. The term ∑*P*_*j*_ stands for the number of public goods consumed by an individual and *b* for the benefit derived from this action. The term ∑*T*_*i*;_ represents the number of consumed toxins. Toxins decrease the overall growth rate and lead to death when they accumulate beyond the threshold value *θ*_*T*_. The parameter θ_*T*_ can thus be understood as a measure of toxin potency. The negative effect toxins have on growth is further defined by the latency parameter κ. If κ = 1.0 then there is no latency and each toxin molecule has a linear additive negative effect on growth rate (Fig. S2). With κ > 1.0, susceptible cells can tolerate toxins at low uptake rates, while the negative effects of toxins on growth accelerate with increased toxin uptake. This accounts for the common phenomenon that bacteria possess non-specific resistant mechanisms (e.g. efflux pumps), thereby often tolerating low toxin concentrations (Fernández and Hancock 2012; Nikaido and Pagès 2012).

The public good producing strains *W* and *P* constitutively secrete one molecule per time step, whereas *P* additionally secretes toxins at the same rate. The diffusion of cells, public goods and toxins (described by the diffusion coefficients *D*_*c*_, *d*_*pg*_ and *d_tox_*, respectively) were modeled according to a Gaussian random walk, with a Gaussian random number generator based on the Box-Muller transform that converts uniformly distributed random numbers to normally distributed random numbers. Following diffusion, public goods and toxins were consumed whenever there was co-localization of molecules and cells.

Both, public goods and toxins were represented by points on the landscape up to the precision of the computer (double precision) (Figs. 1 + S1). The molecules remain in the simulation until they were either consumed or decayed. The probability of decaying was determined by a durability value δ and an exponential decay function

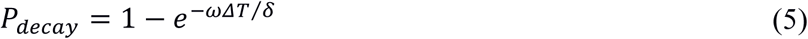

where *ΔT* is the current lifespan of a molecule and *ω* = 0.1 the steepness of the decay.

On our off-lattice landscape, individuals can physically overlap following cell growth and diffusion. To remove the overlaps, we implemented a procedure where cells are moved slightly apart from each other by a random factor scaled by a maximum pulling distance. In a final step, our stimulation involves a life-dead-control of each individual followed by the removal of dead cells that have passed the toxin threshold value *θ_T_*. Fig. 2 depicts the order in which all the actions in our simulation were executed per time step (representing one second). This simulation cycle continues until the total number of cells reaches the carrying capacity of *K* = 1000 cells. For each of the simulated parameter settings, we performed 50 independent replicates. At each time step, we recorded the relative strain frequency. At the end of a simulation, we measured the final strain frequencies, the mean time between two cell divisions, and the *percapita* public good and toxin uptake for each strain.

**Figure 2.**
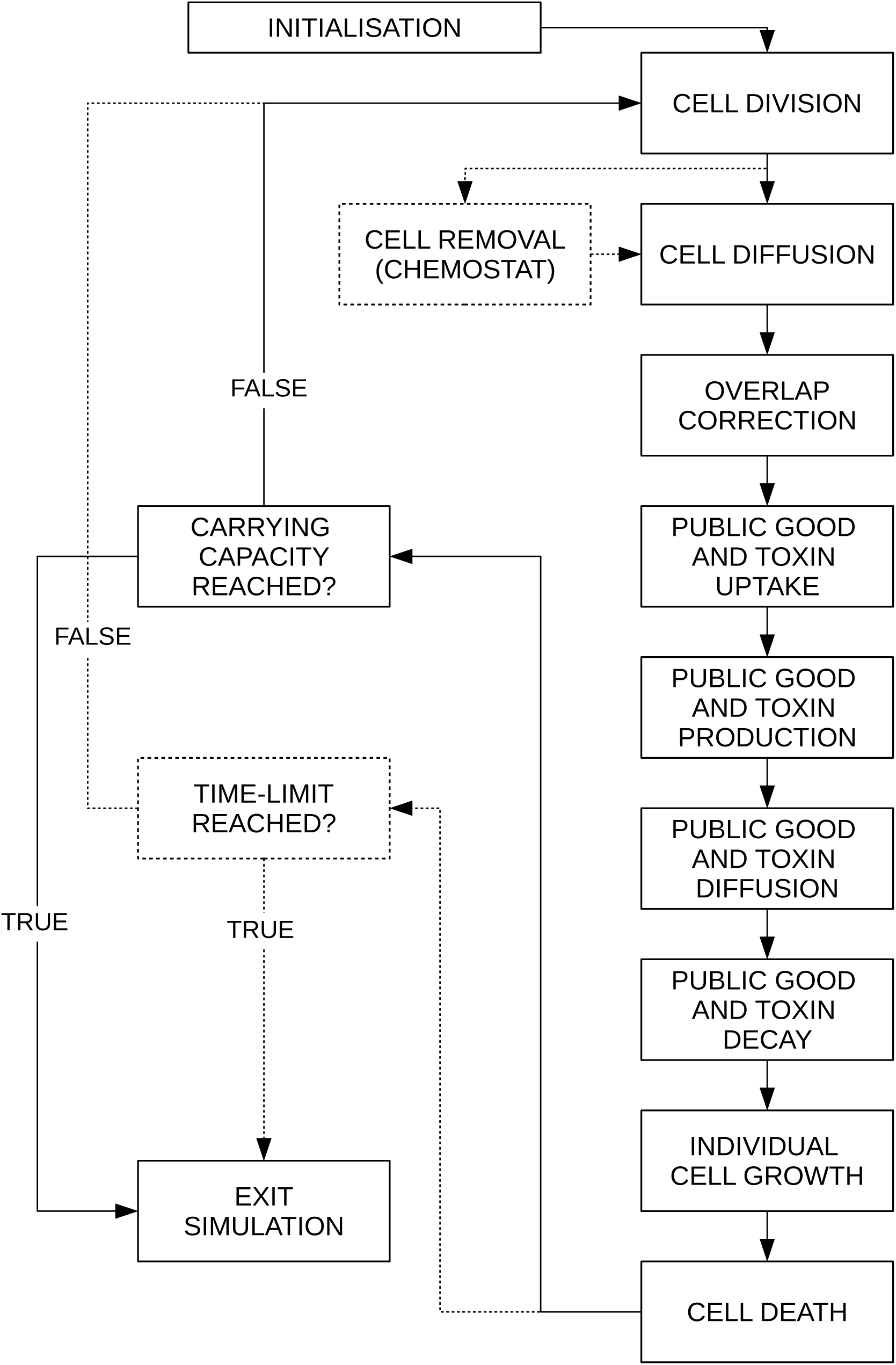
Flow diagram of the computer simulation showing the order of each procedure called during a single time step (i.e. one second). Our platform allows for two different types of culturing conditions: batch-culture growth, where simulations stop when populations reach the carrying capacity *K* = 1000 (solid lines); and continuous growth in a chemostat, where simulations stop after a given time period (dashed lines). To guarantee continuous growth, the chemostat cycle comprised a random cell removal mechanism, such that population size was kept constant at *K*/2. Each box describes a key element in a cell’s life cycle, affecting its fitness: i.e. cells grew according their fitness equations (see main text) and divided when reaching the threshold radius of 1 µm (two times the initial radius). Since cells are free to move on a continuous landscape, they can overlap following diffusion. To remove cell overlap, we used a procedure based on the physics of hard-core interactions used in molecular dynamics and cell elasticity.

### EXPLORED PARAMETER SPACE

Table 1 provides an overview of all the parameters in our simulations and the value ranges explored. One key focus of our study is to understand how cell dispersal and molecule diffusion affect cooperation and policing. For cell dispersal *D*_*c*_, we considered active but random bacterial motility, and varied this parameter from 0 µm^2^/s (no dispersal) to 3.5 µm^2^/s (high dispersal) in steps of 0.5. For molecule diffusion *d*_*pg*_ and *d*_*tox*_, we implemented passive random diffusion processes and varied parameters from 1.0 µm^2^/s (low diffusion) to 7.0 µm^2^/s (high diffusion) in steps of one. Another important aim of our study is to explore how toxin properties affect policing. In addition to toxin diffusion, we varied: (a) toxin durability *δ*_*tox*_ from 50 (rapid decay) to 500 (intermediate decay) to 5000 (low decay) time steps; (b) toxin threshold *θ_T_* from 600 (few molecules required to kill) to 1800 (many molecules required to kill) in steps of 400; and (c) toxin latency κ from 1.0 (low tolerance and immediate linear adverse effect) to 6.0 (high tolerance and delayed exponential adverse effect) in steps of 0.5.

**Table 1.**
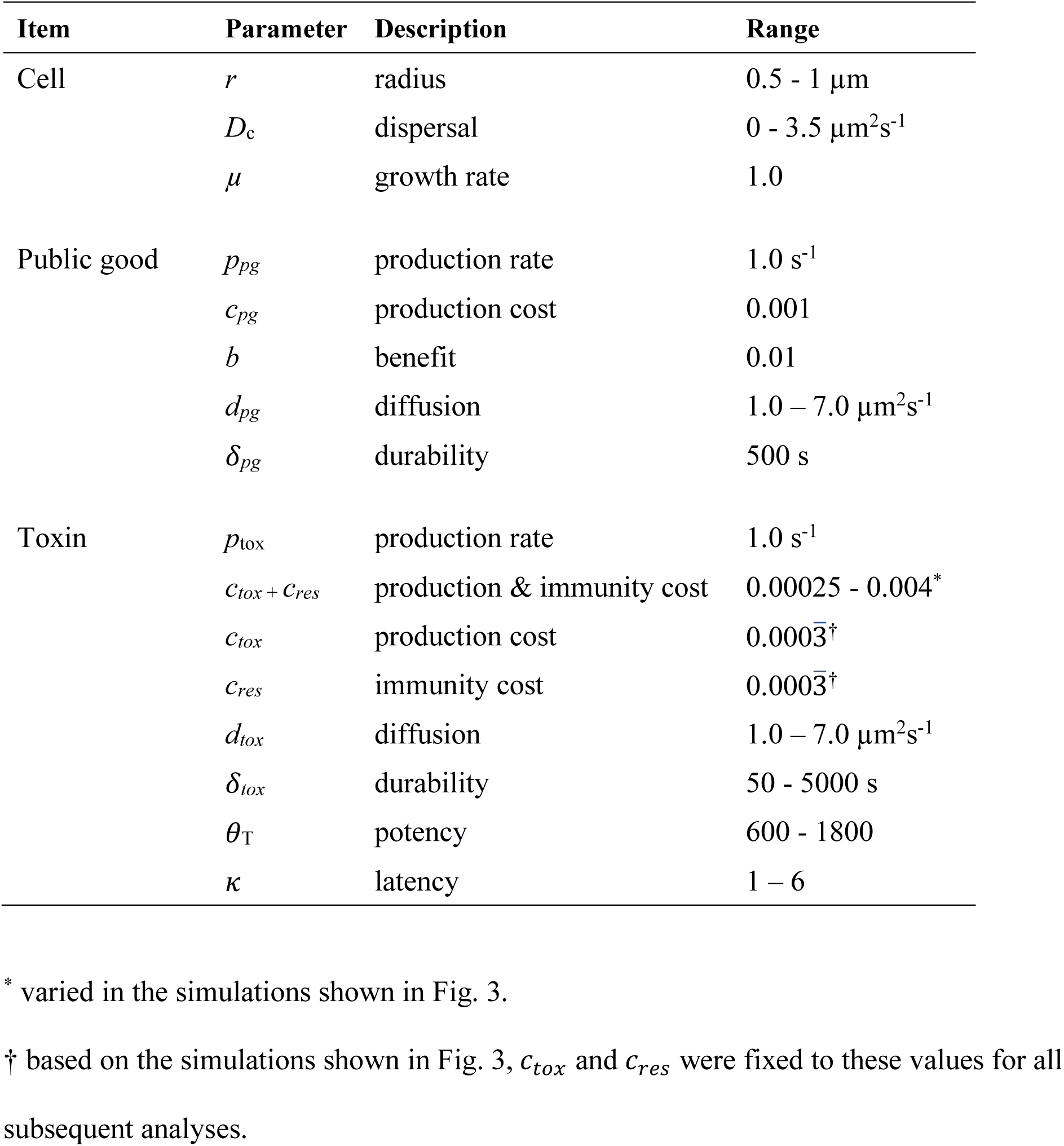

Finally, the outcome of our simulations necessarily depends on the cost and benefit parameters. We implemented a basic growth µ to ensure that all cells can grow even if they do not produce public goods. This ensures that cells can only die because of toxins and not because they experience negative growth. µ = 1.0 results in cell division occurring every 1,200 time steps, in the absence of molecule secretion. We fixed the cost and benefit of public good production to *c*_*pg*_= 0.001 and *b* = 0.01. These values are based on our previous experience with the system (Dobay et al. 2014) and ensure that public production generates a substantial net fitness benefit, thus significantly reducing cell division time (see also Fig. 3). Of key interest is how costly policing (*c*_*tox+res*_) can be relative to *c*_*pg*_ in order to maintain a net benefit of cooperation. To address this question, we gradually varied the cost ratio of these two traits [*c*_*pg*_/(*c*_*tox*_ + *c*_*res*_)] from 0.25 to 4.0. For all simulations, we assumed *c*_*tox*_ = *c*_*res*_.

**Figure 3.**
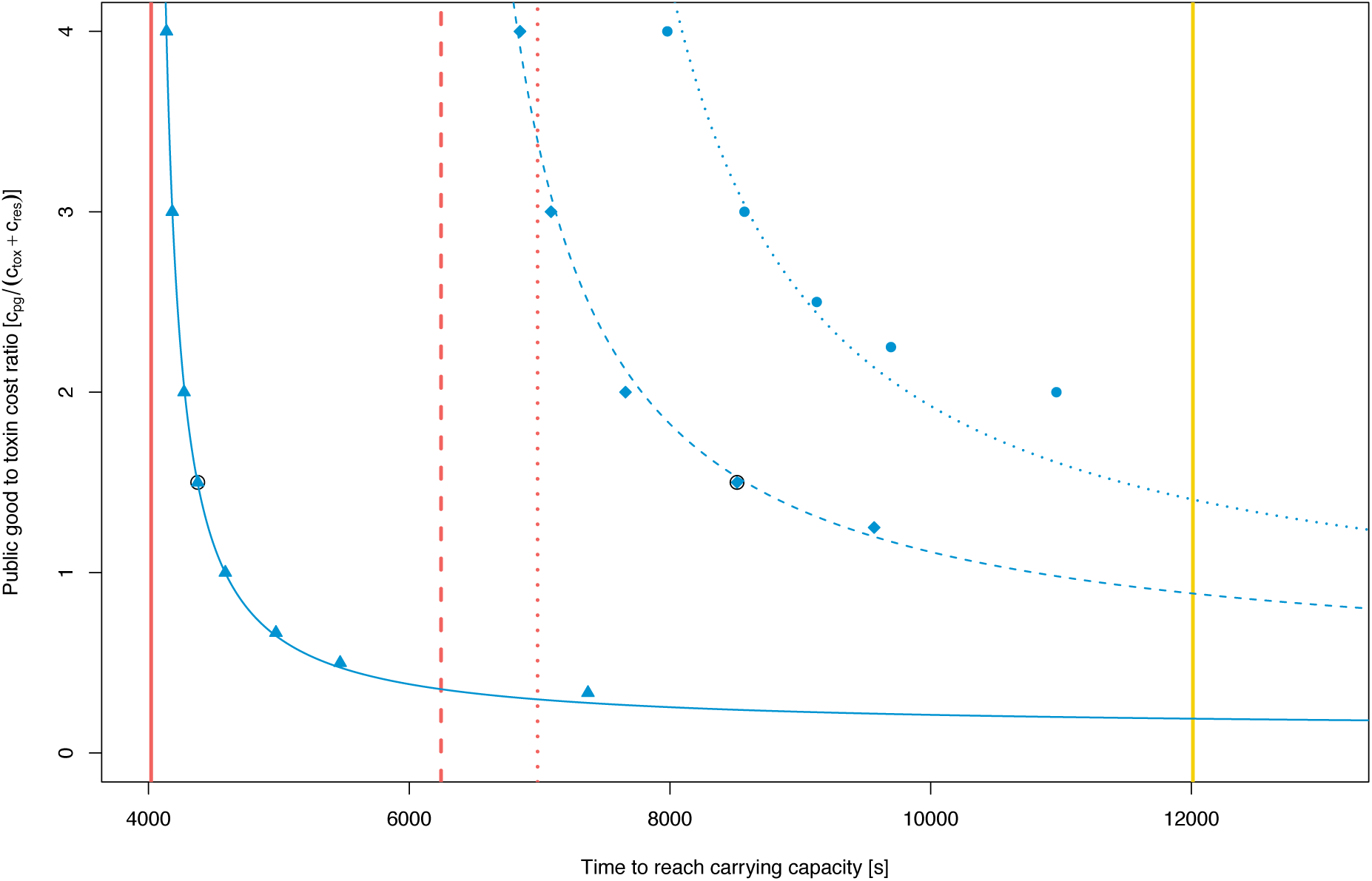
Growth in monoculture and the relative cost of policing. We examined how large the cost of policing (*c*_*tox*_ + *c*_*res*_) can be relative to the cost of public good production (*c*_*pg*_) to guarantee a net benefit of cooperation. To address this question, we compared the growth of the policing strain (producing toxins and public goods, in blue) with the growth of the cooperator strain (producing only public goods, in red) and the cheater strain (producing neither toxins nor public goods, in yellow) in monoculture for a range of cost ratios, under three different diffusion regimes. The yellow bar indicates that the time needed by the cheater strain to reach carrying capacity was not affected by the diffusivity of the environment. The red bars show that the time needed by the cooperator strain to reach carrying capacity increased with higher diffusivity of the environment (plain line: low diffusivity, *D*_c_ = 0.0 µm^2^/s, *d*_pg_ = *d*_tox_ = 1.0 µm^2^/s; dashed line: intermediate diffusivity, *D*_c_ = 2.0 µm^2^/s, *d*_pg_ = *d*_tox_ = 3.5 µm^2^/s; dotted line: high diffusivity, *D*_c_ = 4.0 µm^2^/s, *d*_pg_ = *d*_tox_ = 7.0 µm^2^/s). For policing to evolve, the time needed by the policer strain to reach carrying capacity must be within these boundaries (low diffusivity: blue triangles; intermediate diffusivity: diamonds, high diffusivity: dots). Values represent averages of 50 independent simulations. Standard errors of the mean are smaller than the size of the markers and therefore not shown. We set *c*_*pg*_= 0.001, while varying *c*_*tox*_ + *c*_*res*_ from 0.00025 to 0.004. The black encircled values correspond to *c*_*pg*_/(*c*_*tox*_ + *c*_*res*_) = 1.5, the relative policing cost used for all subsequent simulations.

### STRAIN PERFORMANCE IN MONOCULTURE

In a first set of simulations, we assessed the performance of the strains *W* (wildtype cooperator) *P* (policing cooperator) and *C* (public-good cheater) in monoculture. These assays allow us to quantify (i) the net benefit of cooperation, and (ii) the acceptable boundary costs of policing to ensure a net benefit of cooperation. A net benefit of cooperation is given when the policing strain *P* grows better than strain *C*, which neither produces public goods nor toxins. Because we know that cell dispersal and secreted molecule diffusion influence the efficiency of cooperation (Dobay et al. 2014), we performed simulations under three different diffusivity regimes, including high (*D*_*c*_ = 4.0 µm2/s, *d_pg_* = *d*_*tox*_= 7.0 µm2/s), medium (*D*_*c*_ = 2.0 µm2/s, *d_pg_* = *d*_*tox*_ = 3.5 µm^2^/s) and low (*D*_*c*_ = 0.1 µm^2^/s, *d*_*pg*_ = *d*_*tox*_ = 1.0 µm^2^/s) diffusivity. We then compared the time needed by the three strains to reach carrying capacity.

### PAIRWISE STRAIN COMPETITION

We first simulated competitions between the strains *W* and *C* across the indicated range of dispersal and diffusion parameters to understand the conditions under which cooperation can be maintained in the absence of policing. Next, we competed *P* against *C*. In this scenario, *C* can still exploit the public good produced by *P*, but at the same time is harmed by the toxins secreted by this opponent. In a first set of simulations, we implemented intermediate default toxin properties (*d*_*tox*_ = 4.0 µm^2^/s, *δ*_*tox*_ = 500 s, *θ_T_* = 1000, and κ = 3.5) to examine whether policing extends the range of conditions across which cooperation is favored. In a second set of simulations, we then individually manipulated each of the four different toxin properties to identify the features required for toxin-mediated policing to be efficient.

### THREE-WAY STRAIN INTERACTIONS

To examine what happens when the genetic linkage between cooperation and policing breaks, we performed three-way competitions between strains *P, C*, and *R*. The latter strain no longer produces the toxin, but is resistant to it. Because we do not necessarily expect an overall winner in these competitions, as strain frequency could potentially follow cyclical patterns (Kerr et al. 2002), we extended our simulation code by implementing a random cell removal event, which is activated as soon as more than 30 % of the surface is covered with cells. This allowed us to keep population size at roughly *K*/2, and to follow strain frequency over extended periods of time (i.e. 80,000 time steps).

## Results

### STRAIN PERFORMANCE IN MONOCULTURE

The growth of strain *C* (neither producing public goods nor toxins) was not affected by the diffusivity of the environment, and was solely determined by the basic growth rate µ, reaching carrying capacity after 12,000 time steps (Fig. 3). Strain *W* (producing public goods) grew significantly better than *C*, demonstrating the benefit of public goods cooperation. The growth of this strain was reduced in more diffusive environments, where the likelihood of public good sharing and consumption declines. The performance of strain *P* (producing toxins in addition to public goods) greatly varied both in response to the relative costs of policing and the diffusivity of the environment. In environments characterized by low diffusion, *P* always outperformed *C* even when the cost of toxin production was four times higher than the cost of public good production (triangles in Fig. 3). This was no longer the case in environments characterized by intermediate (diamonds in Fig. 3) or high diffusivity (dots in Fig. 3), where *P* only grew better than *C* when toxins were cheaper than public goods. If this condition was not met, then the high costs of policing combined with reduced public good consumption decreased cell growth to a point that prevented populations from reaching carrying capacity.

Interestingly, the relationship between the cost ratio *c*_*pg*_ /(*c*_*tox*_ + *c*_*res*_) and the time needed to reach carrying capacity (τ_*K*_) was best captured by the Monod equation (Monod 1949), a hyperbolic equation initially used to explain the exponential growth rate as a function of nutrient concentration. Applied to our system, we found that the function

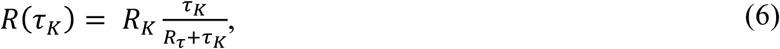

provides a fair approximation to relate *r* to τ_*K*_, where R_*K*_ represents the ratio limit for reaching carrying capacity and R_*τ*_ the time at which the carrying capacity is half the maximum (Fig. S3).

### LOW CELL DISPERSAL FAVORS COOPERATION WITHOUT POLICING

When competing *W* against *C* without a policing mechanism in place, cooperation was only strongly favored when cells did not disperse (Fig. 4A). In other words, if cooperator cells grew as microcolonies, physically separated from the cheaters, then efficient public good sharing among cooperators is promoted. We further found that cooperators and cheaters could coexist when cell and public good diffusion was low, but greater than zero (Fig. 4A). Under all other conditions, cheaters strongly dominated the community and pushed cooperators to very low frequencies. These results are in line with our previous simulations, showing that there is only a narrow parameter space within which public good cooperation is favored (Dobay et al. 2014).

**Figure 4.**
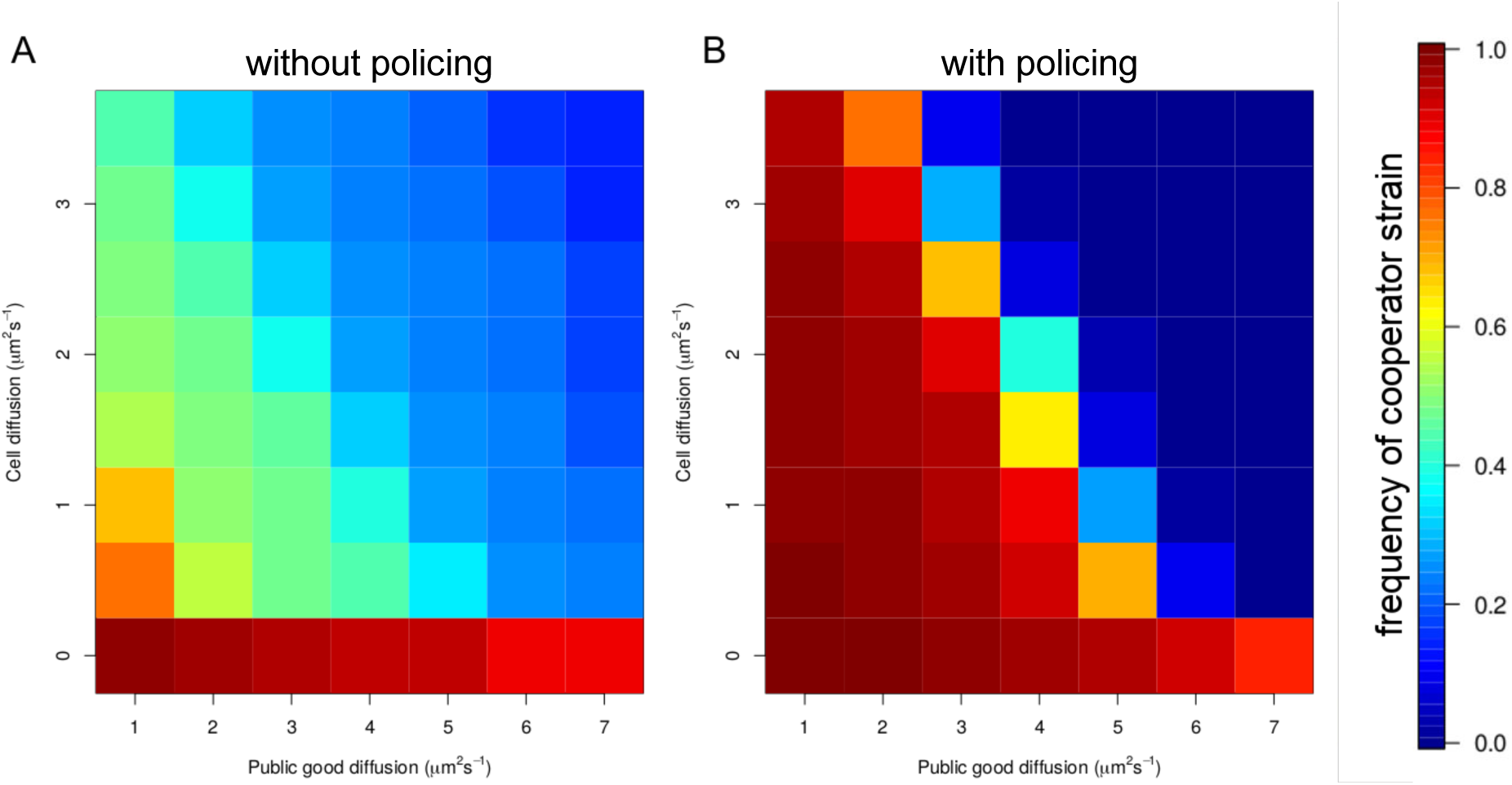
Competition between cheaters and cooperators across a range of cell dispersal *D*_c_ and public good diffusion *d*_pg_ coefficients. Heat maps depict the frequency of the cooperator strain after the community reached stationary phase. (A) Outcome of competitions between the public-good-producing cooperator *W* and the cheater *C* in the absence of a policing mechanism. (B) Outcome of competitions between the policing cooperator *P* and the cheater C. *D*_c_ varied from 0.0 to 3.5 µm^2^/s, whereas *d*_pg_ varied from 1.0 - 7.0 µm^2^/s. The other parameters were set to intermediate values: *d*_tox_ = 4.0 µm^2^/s, *δ*_*tox*_ = 500 s, *θ*_T_ = 1000, Κ = 3.5.

### TOXIN-MEDIATED POLICING HAS POSITIVE AND NEGATIVE EFECTS FOR COOPERATION

The introduction of a policing mechanism, which operates via the secretion of a toxin that selectively targets cheaters, had multiple dramatic effects on the competitive outcome between the cheater *C* and the cooperating policer *P* (Fig. 4B). Policing strongly increased selection for cooperation under conditions where cooperators could previously only coexist with cheaters (compare Fig. 4A and 4B for combinations of low public good diffusion and cell dispersal). This finding demonstrates that policing can indeed extend the parameter space across which cooperation is favored. Conversely, we found that policing also had negative effects and drastically accelerated selection against cooperation, especially under conditions where cheaters previously experienced only a moderate selective advantage (compare Fig. 4A and 4B for combinations of intermediate public good diffusion and cell dispersal). These two opposing effects led to a sharp transition between conditions that either completely favor *P* or *C*, leaving very few conditions where coexistence between the two strains is possible.

### HOW TOXIN PROPERTIES AFFECT POLICING EFFICIENCY

We found that higher toxin diffusion *d*_*tox*_ significantly increased selection for cooperation (Fig. 5A, 5E and 5B), especially under conditions of intermediate public good diffusion and cell dispersal. These results show that policing is particularly efficient when toxins are sent away to target more distant competitors whilst keeping the public good more local for preferential sharing among producers. Our simulations further revealed that high toxin durability *δ*_*tox*_ increases policing efficiency and thus selection for cooperation (Fig. 5C, 5E, 5D). These findings can be explained by the fact that more durable toxins are more likely to reach target cells. Next, we examined the role of toxin potency *θ_T_* (i.e. the number of toxins needed to kill a target cell), and found that high potency is crucial for policing to promote cooperation (Fig. 5F, 5E, 5G). While this finding seems trivial at first sight, the dramatic effects we observed when decreasing toxin potency are remarkable. For instance, a reduction of toxin potency by 45 % (from θ = 1000 to = 1800) already completely negated any benefit of policing, and even increased the parameter space across which cooperation was selected against. Finally, we explored the role of toxin latency κ for policing. Toxin latency is a measure of target cell tolerance: with low values of κ, the fitness of target cells is immediately affected in a linear way, whereas large values of κ mean that target cells can tolerate a certain level of toxins and negative effects only kick in later, but accelerate with higher toxin uptake rates (Figs. S2). We found that low κ values dramatically increased the efiiciency of policing and selection for cooperation (Fig. 5H, 5E, 5I).

**Figure 5.**
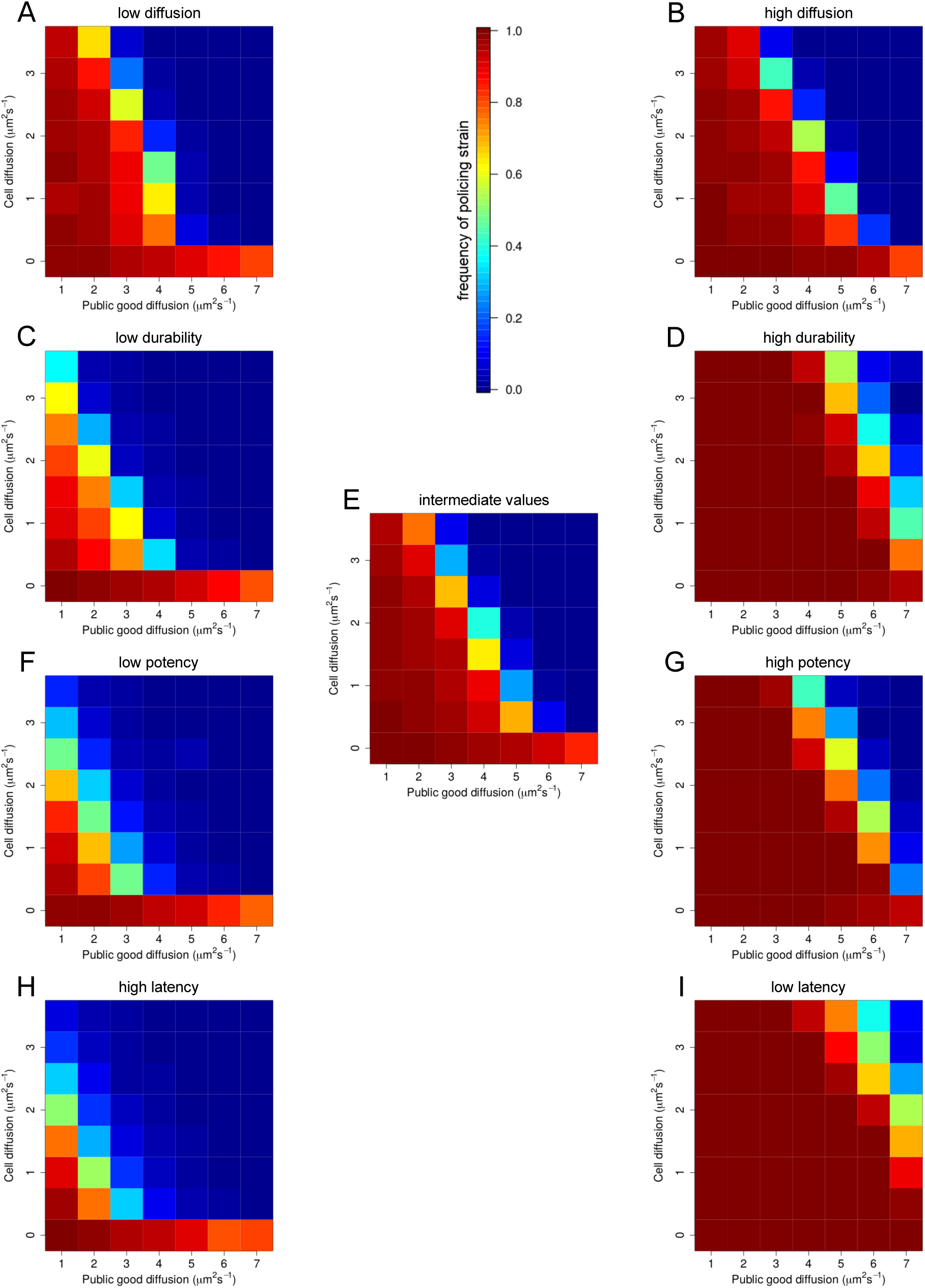
Heat maps exploring the effect of different toxin properties on the success of cooperation. We varied toxin diffusion *d*_*tox*_, toxin durability *δ*_*tox*_, toxin potency *θ_T_* and toxin latency ĸ across a range of values (see Table 1) and show here heat maps of the frequency of the cooperator strain after the community reached carrying capacity for the lowest, highest and intermediate parameter values. At the center of the figure we placed a reference heat map (E), where all the parameters were set to intermediate values (*d*_*tox*_= 4.0 µm^2^/s, *δ*_*tox*_ = 500 s, *θ_T_*= 1000, ĸ = 3.5). We then varied toxin diffusion from 1.0 to 7.0 µm^2^/s (A, E and B), toxin durability from 50 to 5000 (C, E and D), toxin potency from 600 to 1800, (F, E and G) and toxin latency ĸ from 2.0 to 6.0 (H, E and I). We only varied a single parameter at the time, and kept all others at their intermediate default value used for (E).

### POLICING GOES EXTINCT WHEN THE GENETIC LINKAGE BETWEEN TRAITS BREAKS

Next, we simulated the case where the genetic linkage between cooperation and policing breaks. Specifically, we studied the performance of strain *R*, which evolved directly from *P* by losing toxin production but keeping resistance, in competition with *P* and *C* across a range of intermediate environmental diffusivities. We found that the presence of *R* consistently drove *P* to extinction under all environmental conditions tested (Fig. 6). Our simulations, which kept populations at half the carrying capacity (*K*/2 ∼ 500 cells) for an extended period of time (roughly 10 times longer than in the previous pairwise competition assays), allowed us to distinguish three distinct competition phases. The first phase comprises the time frame in which the community grows from three cells to *K*/2. During this phase, we observed cyclical fluctuations of strain frequencies with a general tendency for *C* to increase, *P* to decrease, and *R* to remain stable. The cyclical patterns observed here are reminiscent of the rock-paper-scissors dynamics described by previous studies on bacterial social interactions (Kerr et al. 2002; Kelsic et al. 2015; Inglis et al. 2016), where strains chasing each other in non-transitive interactions in circles with no overall winner. The second phase was characterized by a pronounced dip in *C* frequency accompanied by a strong increase in *R* frequency, and a moderate increase in *P* frequency in eight of the nine diffusion conditions (Fig. 6). This pattern is most likely explained by the accumulation of toxins in the environment, which efficiently suppressed *C*, but at the same time gave *R* leverage, as it could benefit from the effect of policing without paying the cost for it. During the third phase, we observed the extinction of *P*, the concomitant recovery of *C* followed by a decrease of *R* (Fig. 6). These patterns arise because as *P* decreases, toxin concentration declines allowing the recovery of *C*, which then efficiently exploit the public goods produced by *R*. In all cases, our simulations returned to a simple cooperator-cheater scenario, with the relative success of the two strategies being determined by the diffusivity of the environment (similar to the results in Fig. 4A).

**Figure 6.**
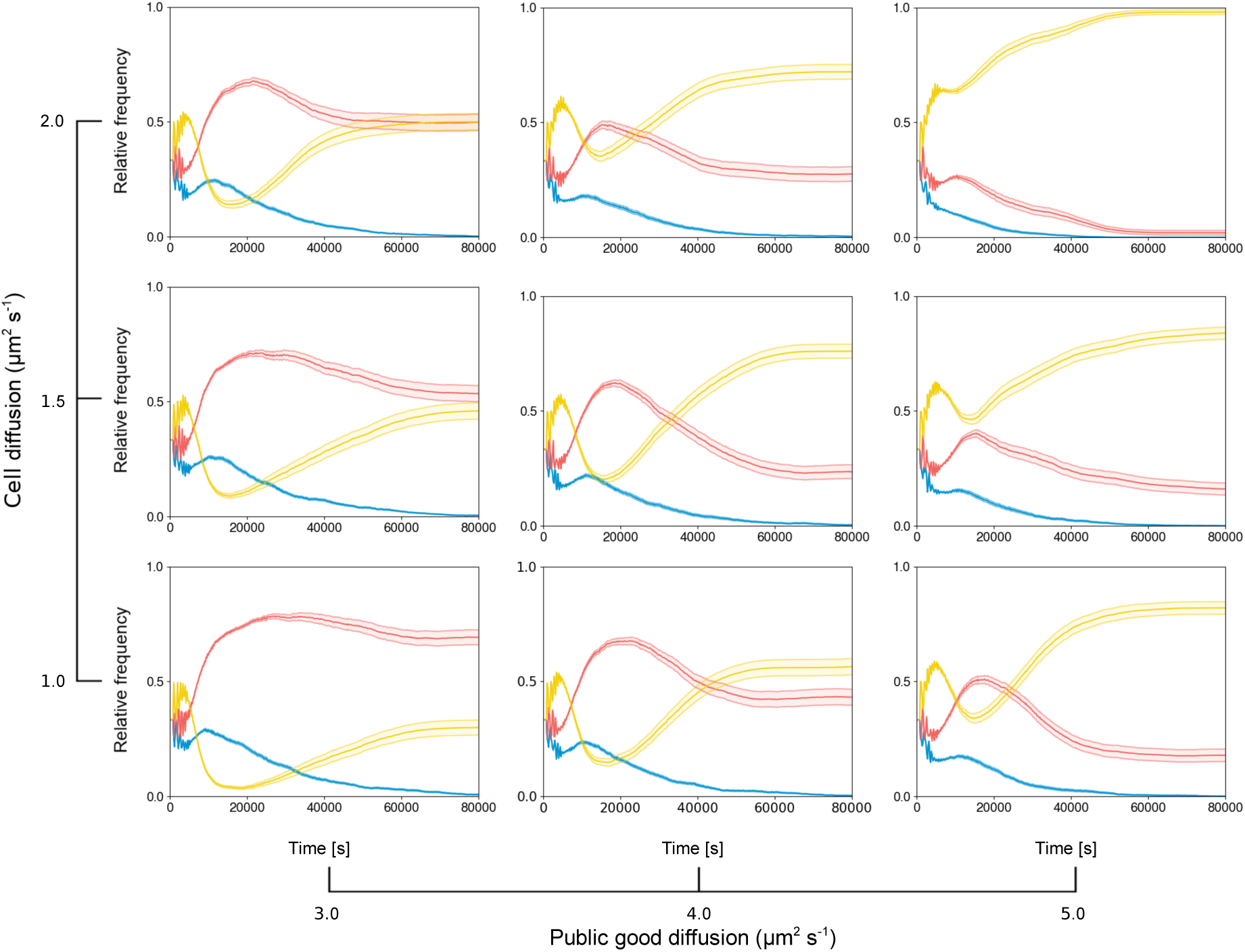
Policing is selected against when the genetic linkage between public good and toxin production breaks. Panels show simulated three-way interactions between cheaters *C* (yellow lines), cooperating policers *P* (blue lines), and toxin-resistant non-policing cooperators *R* (red lines) across a range of public good and cell dispersal values. We assume that *R* can directly evolve from the policing strain *P* when the genetic linkage between public good and toxin production breaks. If this happens, our simulations reveal that policers are always selected against. Trajectories show strain frequency fluctuations in continuous cultures over 80,000 time steps. Values depict means of 50 independent replicates with the standard error of the mean (transparent areas). For all simulations, we kept toxin properties constant: *d*_*tox*_= 7.0 µm^2^/s, *δ*_*tox*_ = 500 s, *θ_T_*= 1000, ĸ = 3.5.

## Discussion

Understanding how policing can repress competition and foster cooperation in social groups of higher organisms has attracted the attention of evolutionary biologists for decades (Clutton-Brock and Parker 1995; Frank 1995, 2003; West et al. 2007b; Ratnieks and Wenseleers 2008; Kümmerli 2011; Raihani et al. 2012; Cant et al. 2013). Here, we explored whether policing could also be an effective way to enforce cooperation in groups of microbes, sharing beneficial public goods. In our models, policing is exerted via the secretion of a toxin that specifically targets cheater cells, which do not contribute to the pool of public goods. Our simulations reveal that policing is most conducive under conditions of intermediate cell dispersal and public good diffusion, where it can extend the parameter space, under which cooperation is favored. We further found that an effective policing mechanism entails a toxin that is: (i) cheaper to produce than the public good; (ii) more diffusible than the public good in order to effectively reach cheaters; (iii) durable; and (iv) potent in killing. However, our simulations also reveal two major downsides of policing. First, policing can accelerate the loss of cooperation under conditions where cheaters and cooperators can coexist in the absence of policing. This leads to a sharp state transition between conditions where policing either favors or disfavors cooperation. Second, policing is selected against if the genetic linkage between public good and the toxin-anti-toxin production can be broken. If this occurs then toxin-resistant mutants that produce public goods, but no longer contribute to toxin production, derail policing, showing that microbial policing itself constitutes an exploitable public good (Inglis et al. 2014).

Several studies have recently proposed toxin-based policing mechanisms to keep cheaters in check (Inglis et al. 2014; Wang et al. 2015; Majerczyk et al. 2016; Evans et al. 2018). While the idea of bacteria being able to punish cheating community members is exciting, our simulations reveal that the ecological spectrum under which policing can be favored is actually quite narrow. For one thing, we found that policing is not required when environmental diffusivity is low, conditions that promote cooperation *per se*. The reason for this is that low public good diffusion and low bacterial dispersal lead to significant spatial segregation of strains within bacterial community, which promotes the local sharing of public goods among cooperators (Kümmerli et al. 2009; Julou et al. 2013; Mitri and Foster 2013; van Gestel et al. 2014; Weigert and Kümmerli 2017). Moreover, we show that policing is not favorable when environmental diffusivity is high, conditions that break any spatial association between cooperators and their public goods. Toxin-mediated policing is detrimental here because: (i) cheaters can freely exploit public goods; (ii) many toxins get lost due to the high diffusion, and thus never reach their targets; (iii) and the high level of cell mixing reduces the efficiency and selectivity of policing, as cooperators are hit by toxins as often as cheaters (Inglis et al. 2011). Overall, it turns out that intermediate levels of environmental diffusivity proved most beneficial for policing. The issue with this relatively narrow parameter space is that environmental diffusivity is likely to vary both temporally and spatially under natural conditions, which could quickly shift the selective balance for or against policing. It thus remains to be seen whether policing can indeed evolve under fluctuating environmental conditions.

Another important point that has received little attention so far concerns the question whether the reported policing mechanisms have indeed evolved for this very purpose or whether they represent by-products of regulatory linkage of traits for other reasons than policing (West et al. 2007c). For instance, in the case of *P. aeruginosa* it is well conceivable that cyanide primarily serves as a broad-spectrum toxin to target inter-specific competitors under high cell density (Bernier et al. 2016). This might be the reason why the expression of cyanide is controlled by quorum sensing, and why it is regulatorily linked to other public good traits, such as protease production, whose expression is also controlled in a density-dependent manner (Darch et al. 2012). Consequently, the observed cyanide-mediated policing exerted by wildtype strains on protease-deficient strains could be a mere by-product of this regulatory linkage. Alternatively, it is also possible that an initial co-incidental regulatory linkage between cyanide and protease later proved useful as a policing mechanism and evolved as such through cooption (Foster 2011). Clearly, further research is needed to uncover the evolutionary history of these putative policing mechanisms, and care must be taken to distinguish between mechanistic (proximate) and evolutionary (ultimate) explanation of observed behavioral patterns (West et al. 2007c).

The potential policing mechanisms reported for microbes and the one implemented in our simulations differ in one important aspect from the policing systems found in higher organisms. Specifically, the difference is that the microbial policing mechanisms are genetically fixed (i.e. strains are either cooperating policers or cheaters), whereas in higher organisms policing is perceived as a conditional strategy, which can be applied to sanction cheaters only if required (Clutton-Brock and Parker 1995; Frank 1995, 2003; Kiers et al. 2003; West et al. 2007b; Ratnieks and Wenseleers 2008; Kümmerli 2011; Raihani et al. 2012). In the latter scenario, individuals can take decisions on whether to cheat or to cooperate, and whether to impose sanctions or not. In certain cases, it was found that the mere threat of policing was sufficient to coerce individuals to cooperate and prevent cheating in the first place (Wenseleers and Ratnieks 2006; Kümmerli 2011; Cant et al. 2013). Microbes clearly lack cognitive abilities for such conditional behaviors, and it is thus not surprising that cheating, cooperating and policing strategies are genetically fixed in these organisms. Nonetheless, we argue that it seems fair to use the term ‘policing’ for the reported behaviors, but also to keep in mind the difference between conditional and fixed strategies.

In summary, our work contributes to the development of an evolutionary concept for policing in microbial systems. It shows how ecological factors, in particular the diffusivity of the environment, interact with the properties of a toxin-mediated policing system, to define the parameter space in which policing can be favored. It further demonstrates how realistic individual-based modelling, tracking both cells and their public goods over time and across space, can be used to deepen our understanding of social interactions in microbes.

## AUTHOR CONTRIBUTIONS

T. W., R. K. and A. D. conceived the study. T. W. and A. D. constructed the model. T. W. performed the simulations. T. W., R. K. and A. D. analyzed the data and wrote the paper.

## ACKNOWLEDGMENTS

We thank Barbara König and Marta Manser for comments, the Services and Support for Science IT (S3IT) at the University of Zurich for technical support with the simulations on the ScienceCloud, and the Swiss National Science Foundation (grant no. PP00P3-139164) and the European Research Council (ERC-CoG grant no. 681295) for funding. The authors declare no conflict of interest with this manuscript.

## SUPPLEMENTARY FIGURES

**Figure S1.**
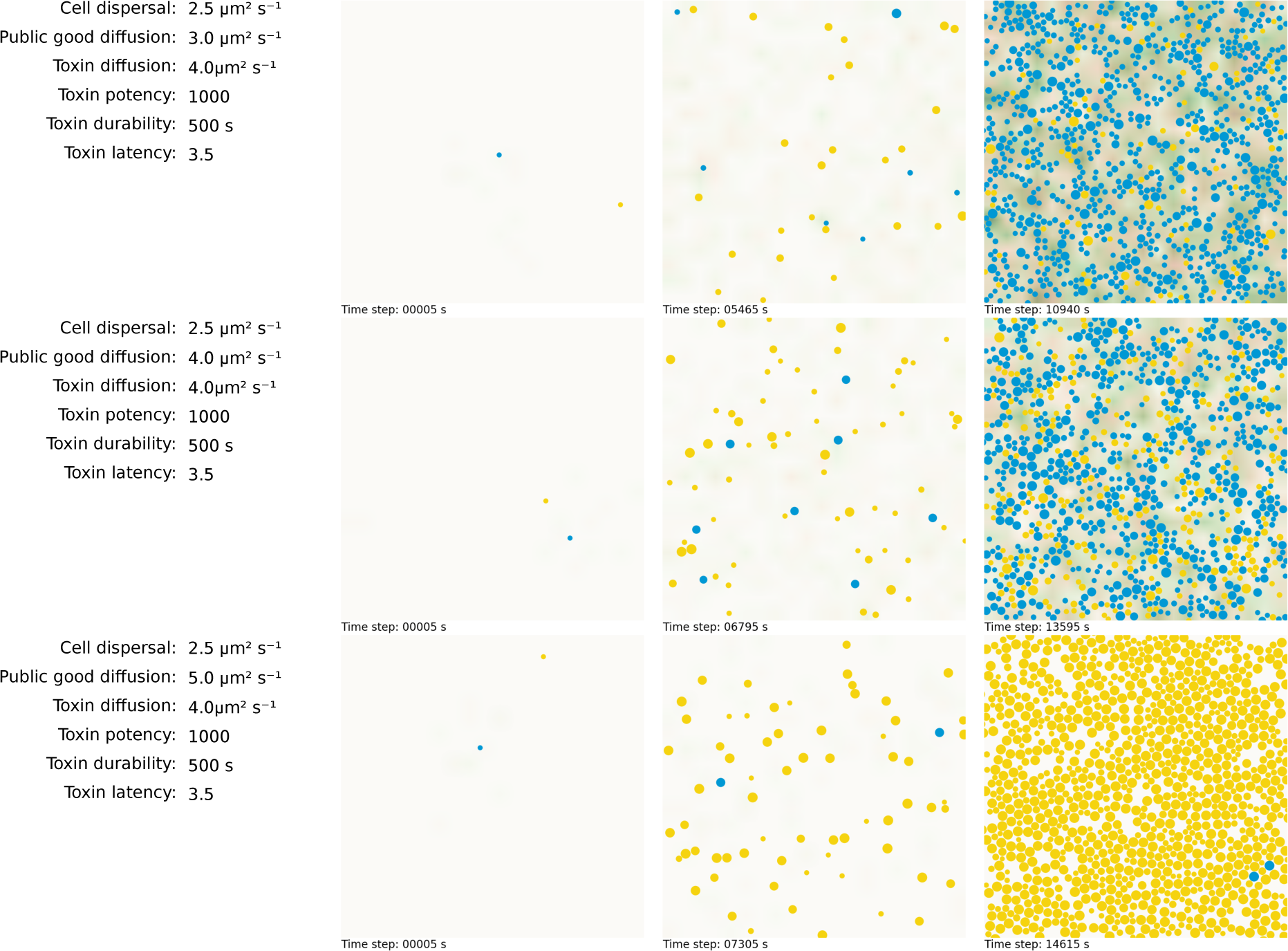
Point-in-time snapshots showing the spatial distribution of cooperating policers (in blue) and cheaters (in yellow) during competition under three different starting conditions. The point-in-time snapshots were taken during the early, intermediate and late simulation phase depicting the growth of digital bacteria in their simulated environments. The only parameter we varied in these examples was public good diffusion *d*_pg_ = 3. The density of public goods and toxins are indicated by the intensity level of the background color. In green are the distribution of public goods and in red the toxins.

**Figure S2.**
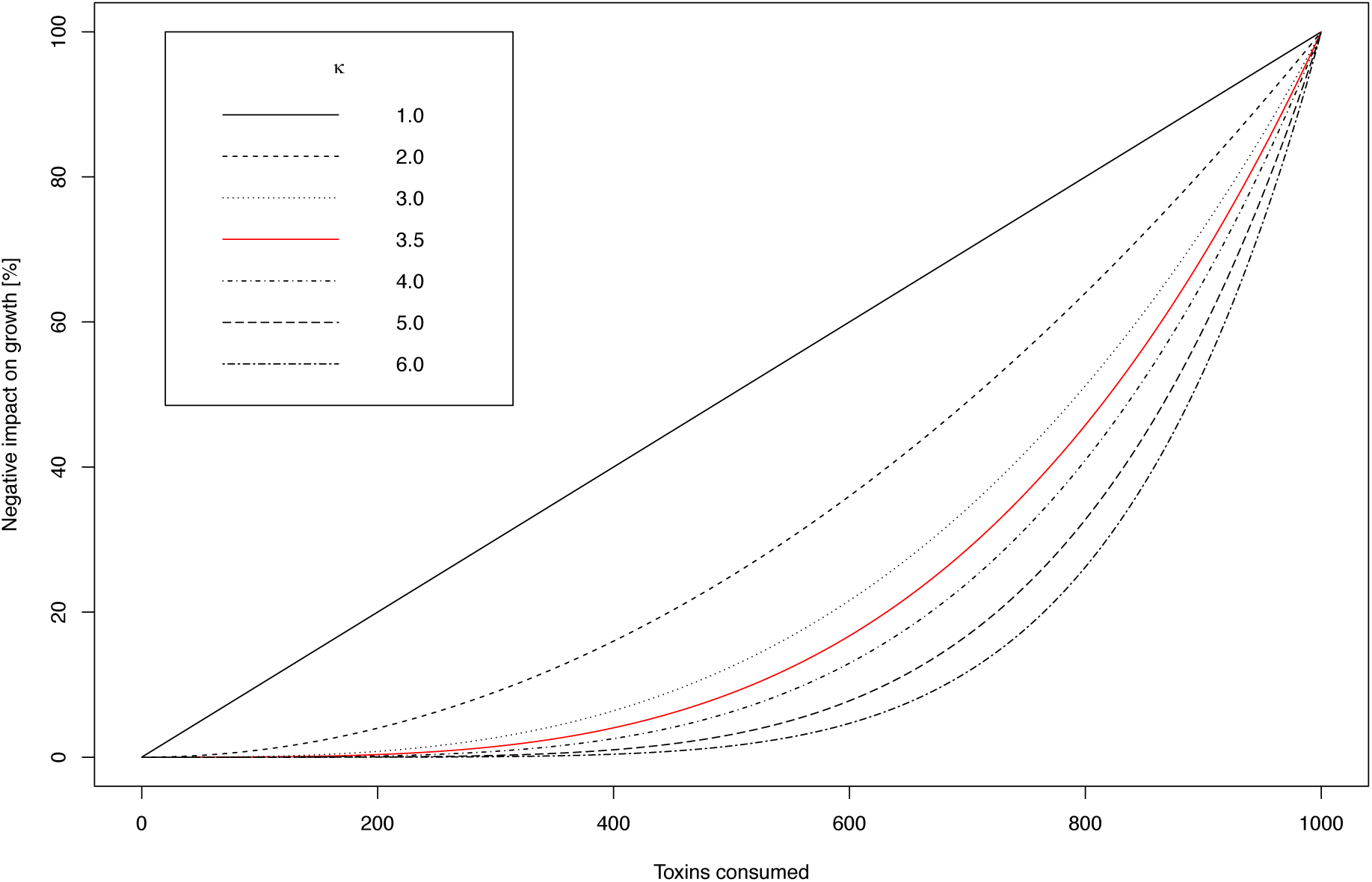
Effect of toxin latency ĸ on the growth rate of sensitive cells. Small values of ĸ lead to an immediate negative impact on growth, whereas larger values of ĸ extend the latency phase during which the growth of sensitive cells is not affected.

**Figure S3.**
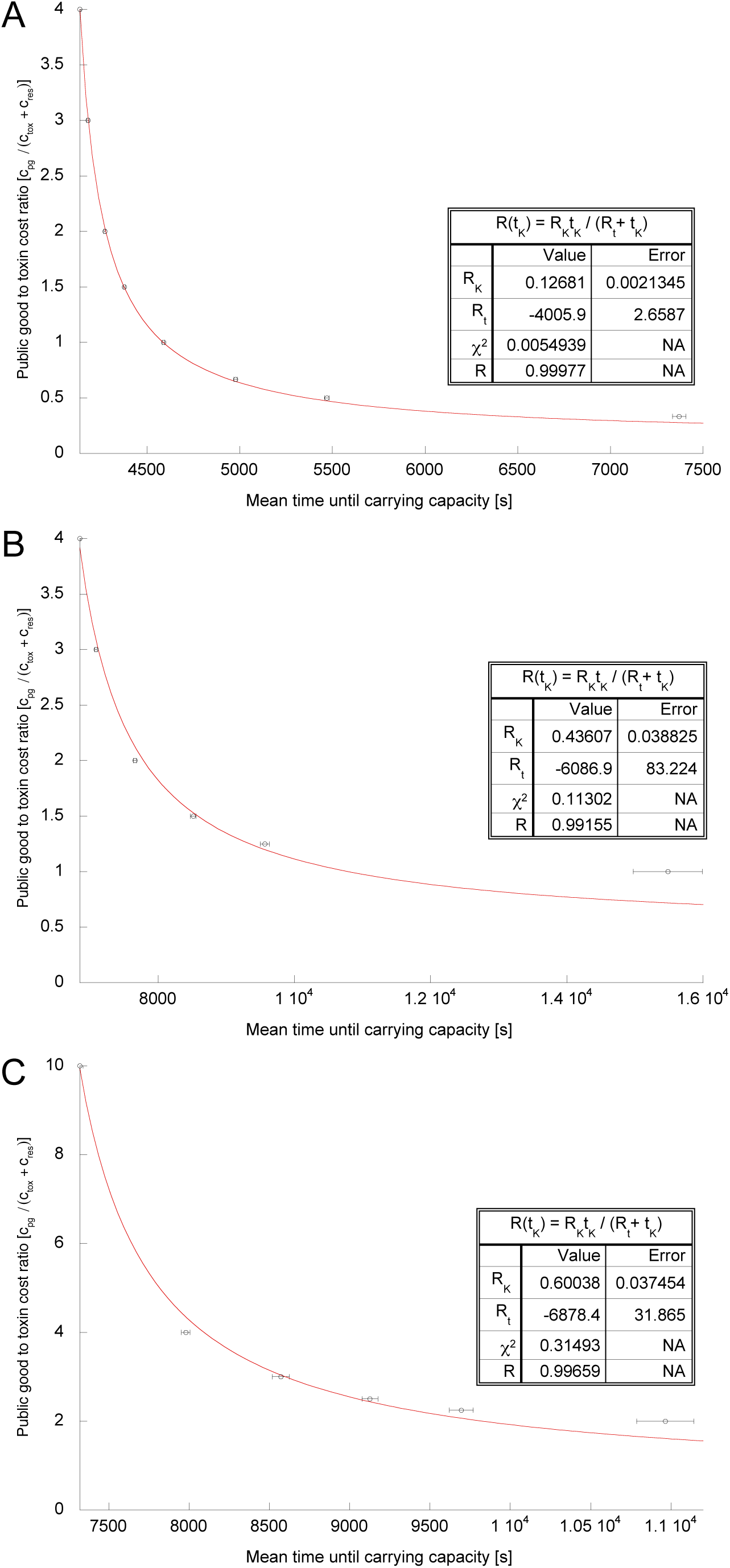
Monod’s equation explains the relationship between the relative cost of policing and strain fitness. We simulated the growth of the policing strain (producing a toxin and a public good) in monoculture for a range of *c*_*pg*_/(*c*_*tox*_ + *c*_*res*_) ratios, under three different diffusion regimes. We kept the cost of public good production constant at *c*_*pg*_ = 0.001, while varying the cost of toxins (*c*_*tox*_ +*c*_*res*_) from 0.00025 to 0.004. The fitness of the policing strain (measured as the time required to reach carrying capacity) declined with higher relative toxin costs. The decline was adequately captured by Monod’s equation under conditions of (A) low diffusion (coefficient of correlation R = 0.999); (B) medium diffusion (R = 0.991); and (C) high diffusion (R = 0.997). However, the equation does not fully capture the observed pattern when the ratio approaches one, namely when (*c*_*tox*_ + *c*_*res*_) ≈ *c*_*pg*_, under conditions of low environmental diffusivity. This is partially understandable since *R*(*τ*_*k*_) does not directly take the viscosity of the medium into account. Details on diffusion parameter values can be obtained from Figure 3. Open circles show means of 50 independent replicates with the standard error of the mean.

